# Map-independent representation of an aggression-promoting social cue in the main olfactory pathway

**DOI:** 10.1101/2021.12.30.474554

**Authors:** Annika Cichy, Adam Dewan, Jingji Zhang, Sarah Kaye, Tiffany Teng, Kassandra Blanchard, Paul Feinstein, Thomas Bozza

## Abstract

While the olfactory system is required for proper social behaviors, the molecular basis for how social cues are detected via the main olfactory pathway of mammals is not well-characterized. Trimethylamine is a volatile, sex-specific odor found in adult male mouse urine that selectively activates main olfactory sensory neurons that express trace amine-associated receptor 5 (TAAR5). Here we show that trimethylamine, acting via TAAR5, elicits state-dependent attraction or aversion in male mice and drives inter-male aggression. Genetic knockout of TAAR5 significantly reduces aggression-related behaviors, while adding trimethylamine augments aggressive behavior towards juvenile males. We further show that transgenic expression of TAAR5 specifically in olfactory sensory neurons rescues aggressive behaviors in knockout mice, despite extensive remapping of TAAR5 projections to the olfactory bulb. Our results identify a specific main olfactory input that detects a prominent male-specific odor to induce inter-male aggression in a mammalian species and reveal that apparently innate behavioral responses are independent of patterned glomerular input to the olfactory bulb.

## Introduction

Species-specific chemical cues strongly influence social behaviors that are critical for survival^1,2^. While many candidate cues (pheromones) have been identified that influence social interactions in vertebrates, few have identified receptors that mediate the observed behavioral effects^3-5^. The olfactory system of mammals comprises two divisions: the main olfactory pathway which responds to myriad volatile, airborne odorants, and the vomeronasal system which primarily detects low volatility molecules—proteins, peptides, bile acids and steroids^3,4,6^. The vomeronasal system exerts a strong influence on behaviors such as maternal and inter-male aggression, sexual receptivity, or reproductive status^7-16^. In a few cases, specific vomeronasal receptors have been identified that drive defined behaviors^16-19^.

While the main olfactory system is thought to serve a generalist function, it has also been implicated in responding to pheromones and social cues^20-27^. However, very little is known about the ligands and receptors involved (see reference 28). The main olfactory pathway uses three classes of G-protein coupled receptors to detect odorants: canonical class I and class II odorant receptors (ORs) and a smaller family of Trace Amine Associated Receptors (TAARs)^29-31^. These three receptor classes are mapped to distinct domains of the olfactory bulb^32-34^, the first relay for olfactory information in the brain, and these may relate to different innate and learned behaviors^34-36^.

Trace Amine Associated Receptor 5 (TAAR5) is expressed in main olfactory sensory neurons and is the most sensitive receptor for the volatile odorant trimethylamine (TMA)^37^. TMA is a sex- and species-specific odor cue that is enriched in the urine of adult male (but not juvenile or female) *Mus musculus*^38,39^. While it has been reported to elicit attraction in mice^39-41^, how TMA influences social interactions is not known. Here we show that TMA, acting through TAAR5, promotes inter-male aggression, revealing a single, defined main olfactory input that drives a key social behavior in mammals.

## Results

### Trimethylamine valence is modulated by social status

Our previous work had shown that amines, such as phenylethylamine, elicit aversion in naïve (untrained) mice via the TAARs^42^. During the course of these experiments, we observed variability in the behavior of group-housed mice towards TMA. We reasoned that behavioral responses to TMA through TAAR5 may depend on factors such as social status (*i*.*e*. dominant vs. subordinate). To test this, we measured valence responses to TMA in male mice of defined social status using a two-chamber place preference assay. These experiments were done in TAAR5 knockout mice (ΔT5) and wild-type littermate controls to determine whether any observed valence effects are receptor-dependent (Fig 1a). Adult male mice were singly-housed with females for a week, then pair-housed with an adult male of the same genotype for one week, after which their dominant/subordinate status was determined based on differential scent marking^43^. Dominant and subordinate males were then tested in the place preference assay (Fig 1b) and preference index (PI) calculated (negative values indicating aversion and positive values indicating attraction).

**Figure 1.**
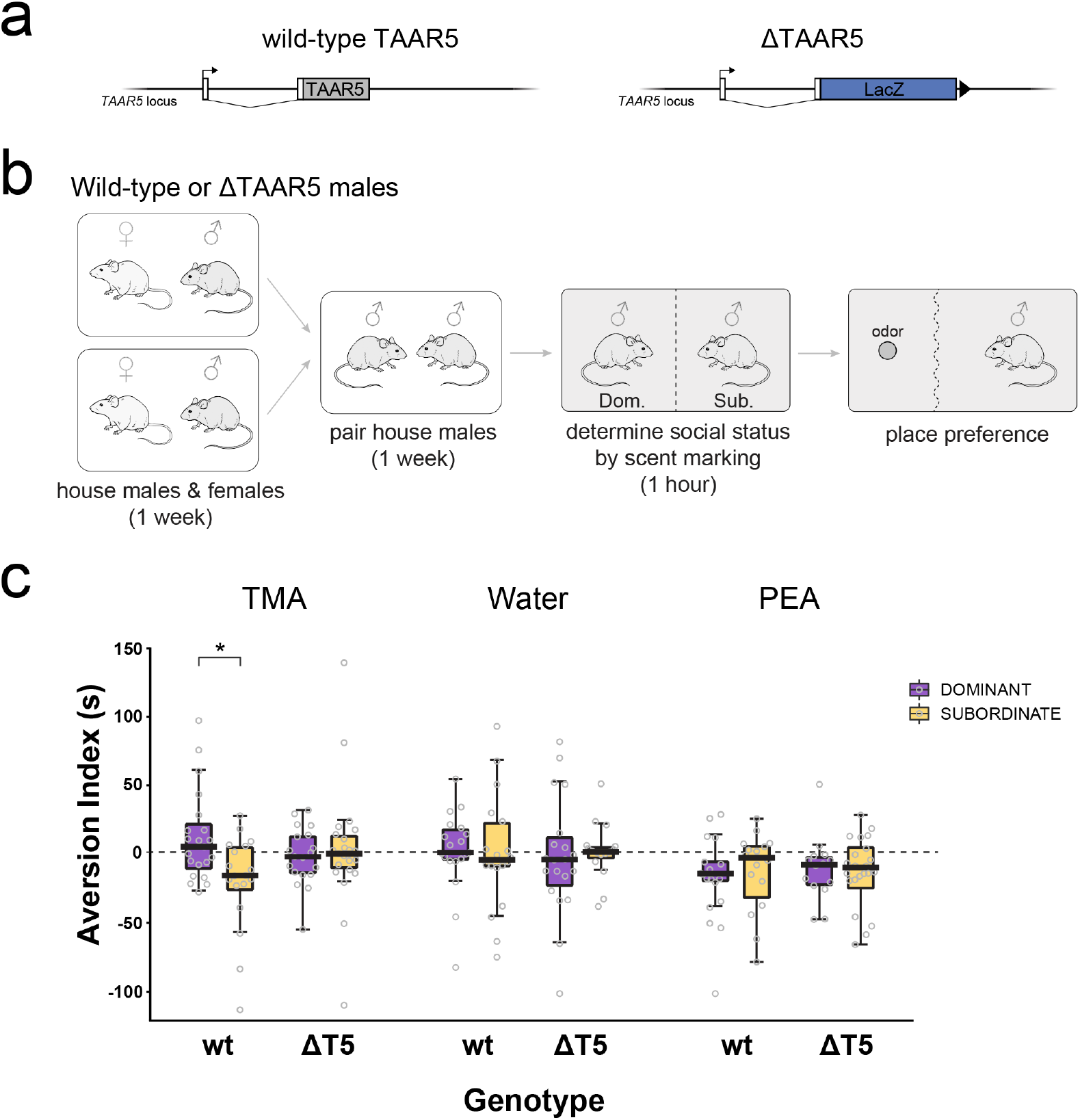
TAAR5 and state dependent valence of trimethylamine. **a)** Top: diagram of wild-type and targeted-deletion TAAR5 alleles. Non-coding exons (white) and the coding sequences for wild-type TAAR5 (grey) and β-galactosidase insertion (blue) are indicated. **b)** Wild-type or ΔTAAR5 males were housed with wild-type females, then pair-housed with a male of the same genotype to establish social dominance. The social status of each male was assessed, and each individual was then tested for odor valence in a place preference assay. Place preference was measured in a clean cage divided into an odorized and non-odorized compartments by a flexible curtain (dotted wavy line). Odor stimuli were delivered in a perforated dish and occupancy time was recorded via video tracking. **c)** Place preference data for dominant and subordinate males of two genotypes, wild-type and ΔTAAR5 to 2% trimethylamine (TMA), water and 10% phenylethylamine (PEA) are plotted as Aversion Index (thick lines, medians; boxes, 25th–75th percentiles; whiskers, most extreme data points not considered outliers using method of Tukey). Index is the difference in occupancy time for a given stimulus and water control (value is positive for attraction and negative for aversion). n = 14-23 mice per experimental group. * p=0.005, ANOVA contrasts (see methods).

On average, males avoided the known aversive odorant phenylethylamine^42,44^ (p<0.001), but showed no preference for the negative control stimulus, water (p<0.643, t-test for mean PI different from 0; Fig 1c). There was no difference in PEA aversion across genotype and state (univariate ANOVA, genotype-by-state interaction, p=0.645). In contrast, male responses to TMA differed according to genotype and social status (univariate ANOVA, genotype-by-state interaction, p=0.024; ANOVA contrasts p=0.005). Wild-type subordinate males displayed aversion, while wild-type dominant males showed mild attraction (p<0.001, t-test for mean PI different from 0). Moreover, the differential response to TMA was abolished in ΔT5 mice (Fig 1c)—no significant difference was observed between ΔT5 dominant and ΔT5 subordinate mice (p = 0.727, ANOVA contrasts). These data indicated that male mice exhibit state-dependent responses to the putative social cue TMA, and that the responses depend on the function of TAAR5.

### TAAR5 influences inter-male aggression

Given the effect of social status on TAAR5-mediated TMA valence, we reasoned that TAAR5 may play a role in male-male social interactions. Consequently, we tested whether TAAR5 influences male interactions during the establishment of dominance. To do this, adult wild-type or ΔT5 males were singly housed with females for a week, and subsequently placed with a male of the same genotype in a clean cage, where the first 30 minutes of behavioral interactions were recorded to digital video (Fig 2a). After this initial interaction, the males were housed together for one week, after which social status was determined via differential scent marking. Videos were scored for a set of 12 behaviors: self-grooming, chasing, anogenital investigation, facial investigation, freezing, mounting, defensive upright, fighting, biting, tail-rattling, pinning, shoveling (Fig S1).

**Figure 2.**
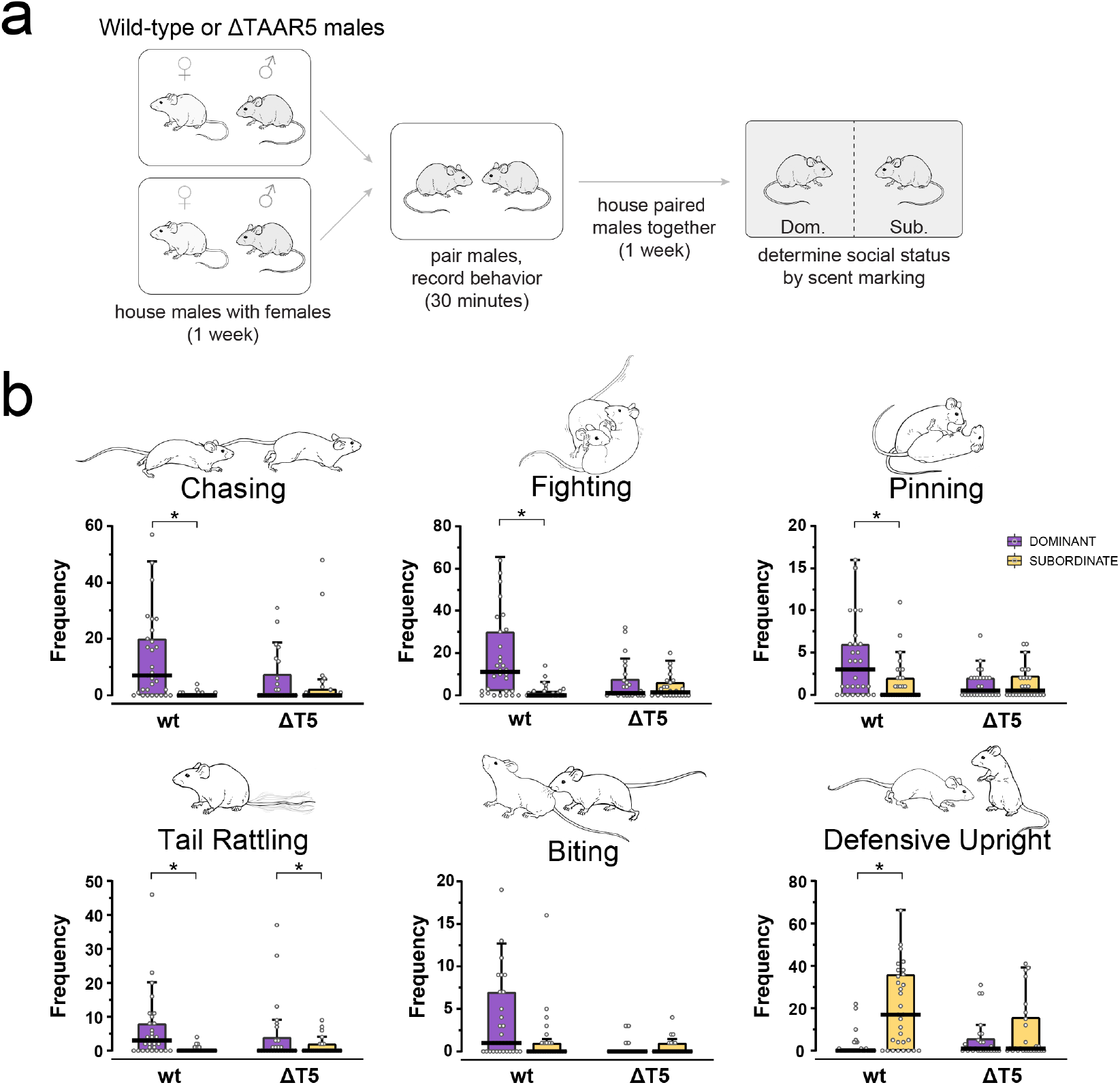
TAAR5 deletion reduces aggression. **a)** Male wild-type and ΔTAAR5 littermates were housed with wild-type females for a week, then pair-housed in a new cage with a male of the same genotype. The first 30 minutes of the interaction was video recorded for quantification of 12 behaviors (Fig S1). Males were then housed together for 1 week, and social dominance assessed using a scent marking assay. **b)** Frequencies of aggression-related behaviors (chasing, fighting, pinning, tail-rattling, biting and defensive upright stance) graphed by genotype (wild-type vs. ΔTAAR5) and social status (dominant, purple vs. subordinate, yellow) of the male initiating the behavior. (Thick lines, medians; boxes, 25th–75th percentiles; whiskers, most extreme data points not considered outliers, method of Tukey). n = 26-31 mice per experimental group. \**p* < 0.0001, chasing, fighting, tail-rattling, defensive upright; \**p* = 0.0072, pinning (negative binomial regression).

We observed a pronounced effect of both social status and genotype on aggressive behavior. Dominant mice are more aggressive than subordinates^45^. Consistent with this, wild-type dominant males showed a significantly higher frequency of chasing (*p* < 0.0001), fighting (fight initiation) (*p* < 0.0001), pinning (*p* = 0.0072), and tail-rattling (*p* < 0.0001; negative binomial regression) as compared with wild-type subordinate mice (Fig 2b). In contrast, ΔT5 mice exhibited no difference in these aggressive behaviors between dominant and subordinate mice (*p* = 0.7543, *p* = 0.307, *p* = 0.7829, respectively) except for tail rattling (*p* = 0.0237; negative binomial regression; Fig 2b). Biting frequency followed the same trend (Fig 2b), but the genotype-by-status interaction was not statistically significant. In addition, we observed differences in aggression-related behaviors by social status and genotype. Subordinate mice typically show defensive behaviors in response to aggression^45^. Consistent with this, wild-type subordinates exhibited a higher frequency of defensive upright posturing as compared with wild-type dominant males (*p* < 0.0001; negative binomial regression; Fig 2b). This difference was abolished in ΔT5 mice (*p* = 0.3238).

The effects of deleting TAAR5 were selective for aggression-related behaviors, as no differences were observed in the frequency of anogenital or facial investigation, freezing, mounting, or shoveling (negative binomial regression, genotype-by-status interaction, *p* > 0.1; Fig S2). We did note an increase in grooming in subordinate vs dominant (wild-type) mice (*p* = 0.0001). This behavior is neither aggressive nor defensive but may be elevated in subordinate mice due to wounding. Consistent with this interpretation, the difference in grooming in dominant vs. subordinates was abolished in ΔT5 mice (*p* = 0.858, negative binomial regression; Fig S2).

Strikingly, deletion of TAAR5 significantly reduced the incidence of aggression-induced mortality. Extended pair-housing to establish dominance occasionally resulted in the death of the subordinate male. The proportion of pairings that resulted in mortality was significantly reduced in ΔT5 mice compared with wild-types: 18 mortalities in 99 pairs (18%) for wild-type vs. 5 mortalities in 86 pairs (6%) for ΔT5 (Fisher Exact Test, *p* = 0.013).

Despite the marked reduction in aggression, TAAR5 deletion males establish a social hierarchy. We found no difference between the proportion of wild-type and ΔT5 mice that exhibited clear differential scent marking: 80 pairs out of 118 (68%) for wild-type vs. 70 pairs out of 105 (67%) for ΔT5 (Fisher Exact Test, *p* = 0.89).

Taken together, our data demonstrate that deletion of the TMA receptor TAAR5 causes a pronounced reduction in inter-male aggression and related defensive behaviors.

### TMA induces inter-male aggression

Next, we tested whether TAAR5-mediated aggression is enhanced by the preferred TAAR5 ligand TMA. Males do not have high concentrations of TMA in their urine until postnatal day 21^39^. Therefore, we swabbed juvenile (P18) wild-type males with a small volume of either water or TMA (diluted in water), paired them with individually-housed adult males and scored aggressive behaviors for the 30 minutes. Since mice do not exhibit adult aggression or defensive responses until >P30 ^46^, we expected instances of reciprocally aggressive behaviors (e.g. fighting, chasing, defensive upright) would be low, and that unilateral actions of the adult male (e.g. biting) would be more informative. As predicted, instances of fighting and chasing were relatively low and not different in response to painted and non-painted subjects (fighting, *p* = 0.4976; chasing, *p* = 0.4755; Fig S3). In contrast, we observed a significant increase in the frequency of biting (negative binomial regression; *p* = 0.0254) in response to TMA-painted as compared with the water-painted juveniles (Fig 3). Thus, the presence of TMA enhanced the level of aggression exhibited towards juvenile male mice. Interestingly, the adult males showed mounting behavior, which was significantly reduced with TMA-painted juveniles (negative binomial regression; *p* = 0.0217). Thus, it is possible that TMA has an effect on sexual behavior in this context.

**Figure 3.**
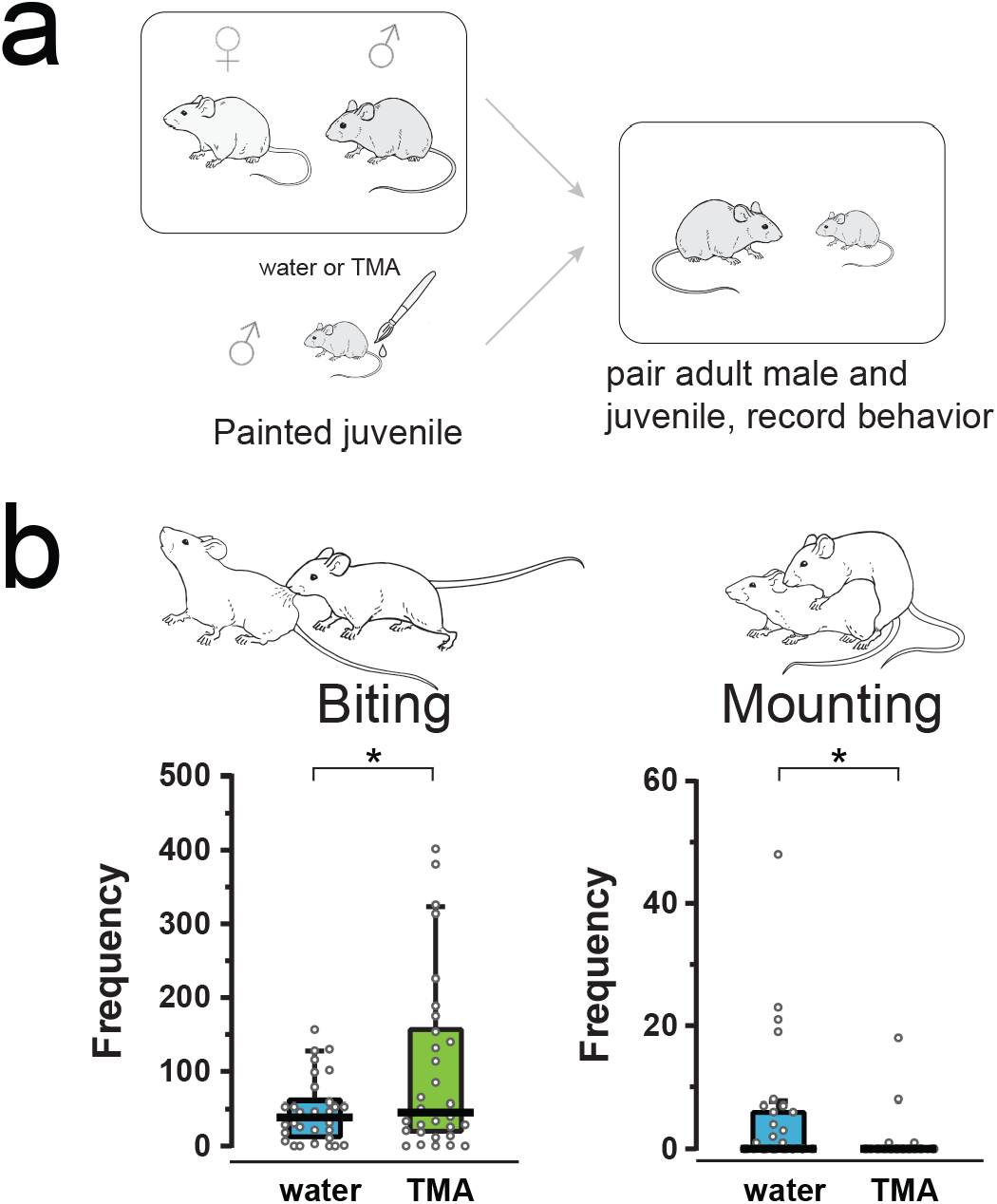
TMA enhances aggression towards juvenile males. **a)** Adult male wild-type mice were housed with wild-type females for a week, then paired in a new cage with a wild-type juvenile (P18) which was painted with water or 5 mM trimethylamine (TMA) in the urogenital region. The first 30 minutes of the interaction was video recorded for quantification of 12 behaviors. **b)** Frequencies biting and mounting exhibited by adult males towards juveniles painted with water (blue) or trimethylamine (TMA, green). Thick lines, medians; boxes, 25th–75th percentiles; whiskers, most extreme data points not considered outliers, method of Tukey). \**p* = 0.0254, biting; \**p* = 0.0217, mounting. n = 30 pairs per experimental group.

### TAAR5 affects social behavior via the main olfactory system

Because the ΔT5 allele is a constitutive knockout, behavioral effects could be due to TAAR5 function in the brain^47^. We were unable to detect expression of the TAAR5 locus in the brain using the ΔT5 allele (which encodes β-galactosidase under control of the endogenous *Taar5* promoter; Fig S4a), and observed variable expression using quantitative RT-PCR in some hypothalamus and amygdala samples (Fig S4b), potentially indicating low levels of expression^47^.

To rule out a central role for *Taar5* in aggression, we asked whether the loss of aggressive behaviors in ΔT5 mice could be rescued by olfactory-specific TAAR5 expression. We generated a transgenic mouse line, *T5-RFPtg*, in which TAAR5 (along with a red fluorescent protein marker) is expressed in a subset of olfactory sensory neurons using a modified OR promoter (Fig 4a). In this strain of mice (T5tg), TAAR5-expressing sensory neurons are located predominantly in the dorsal olfactory epithelium, and send axons to multiple glomeruli in the dorsal olfactory bulb (Fig 4a,b). No detectable RFP expression was observed in other olfactory subsystems (*i*.*e*. vomeronasal organ or Grüneberg ganglion). Moreover, transgene expression could be detected reliably by qPCR in the olfactory bulb but not in brainstem, amygdala, cortex, or hypothalamus (Fig S4b), consistent with olfactory-specific expression of OR promoters/enhancers.

**Figure 4.**
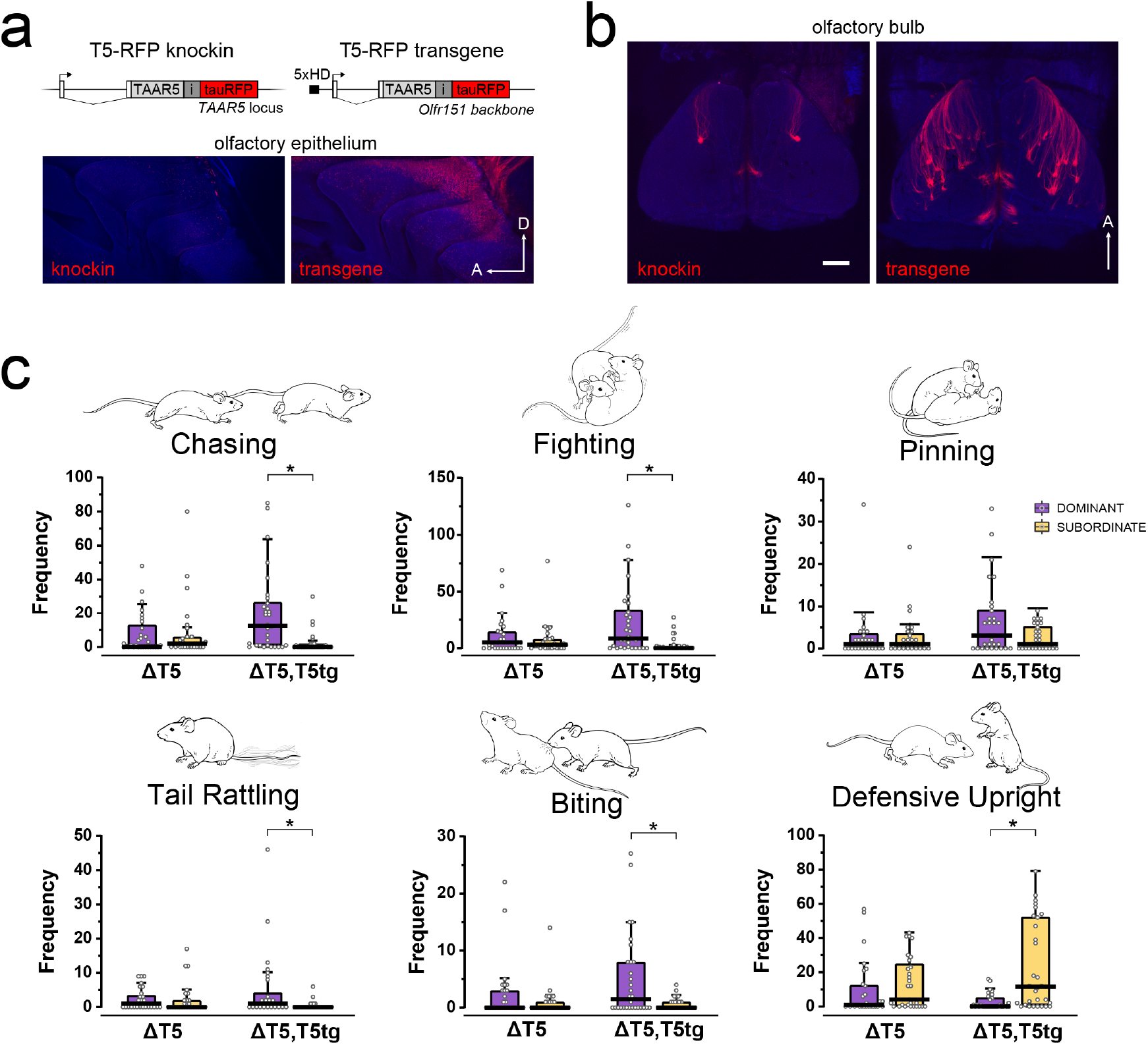
Transgenic expression of TAAR5 in the olfactory system rescues aggression in ΔTAAR5 mice. **a)** Top, diagram of the targeted TAAR5 locus and the TAAR5 transgene (T5-RFPtg). Non-coding exons are shown (white boxes). Coding sequences for wild-type TAAR5 (grey) and tauRFP reporter (red) are separated by an internal ribosome entry site (“i”, dark grey) as indicated. The transgene backbone is a fragment of the *olfr151* locus with a 5x repeat homeodomain sequence (5xHD; black box) inserted upstream of the transcription start site (arrow). Bottom, confocal images showing expression of TAAR5 (RFP; red) in wholemounts of the olfactory epithelium. **b)** Confocal images of dorsal olfactory bulb showing axonal targeting to glomeruli. Scale bar = 500 μm. **c)** Frequencies of aggression-related behaviors (chasing, fighting, pinning, tail-rattling, biting and defensive upright stance) graphed by genotype (wild-type vs. ΔTAAR5) and social status (dominant, purple vs. subordinate, yellow) of the male initiating the behavior. (Thick lines, medians; boxes, 25th–75th percentiles; whiskers, most extreme data points not considered outliers, method of Tukey). n = 29-30 mice per experimental group. \**p* < 0.0001, fighting, tail-rattling, defensive upright; \**p* = 0.0001, chasing, biting (negative binomial regression).

Transgenic expression of TAAR5 rescued differences in aggressive behaviors between dominant vs. subordinate ΔT5 mice (Fig 4c). Consistent with our previous experiments (above), ΔT5 dominant and subordinate males showed equivalent amounts of fighting (*p* = 0.4348) and chasing (*p* = 0.9253; negative binomial regression; Fig 4c). In contrast, transgene-containing deletion Taar5 (ΔT5;T5tg^+^) dominant males exhibited significantly elevated fighting (*p* < 0.0001), chasing (*p* = 0.0001), biting (*p* = 0.0001), and tail-rattling (*p* < 0.0001, negative binomial regression) as compared with ΔT5;Tg^+^ subordinate males (Fig 4c). Similarly, the transgene also rescued defensive behaviors in subordinate mice. The frequency of defensive upright posturing was equivalent in ΔT5 dominant and subordinate mice (*p* = 0.4508), but increased significantly in ΔT5;T5tg^+^ subordinates (*p* < 0.0001; Fig 4c). We also noted a significant increase in anogenital investigation initiated by dominant ΔT5;T5tg^+^ males (*p* = 0.0011, negative binomial regression; Fig S5).

Notably, the proportion of male-male pairings that resulted in mortality was also rescued in ΔT5;T5tg^+^ mice as compared with ΔT5: 16 mortalities in 112 pairs (14%) for ΔT5;T5tg^+^ vs. 5 mortalities in 101 pairs (5%) for ΔT5 (*p* = 0.0362; Fisher’s Exact Test). Thus, the effect of TAAR5 deletion on inter-male aggression is mediated by the main olfactory system.

### TAAR5-mediated aggression is map-independent

It has been suggested that regions of the olfactory bulb are dedicated to learned or innate odor responses^34,35^, raising the possibility that TAAR5-mediated aggression requires wiring to specific glomeruli. We note that the *T5-RFPtg* line dramatically remaps TAAR5 inputs in the olfactory bulb. The transgene is expressed in 10.3-fold higher number of OSNs than the native TAAR5 knock-in (mean cell counts in matched sections: 223 ± 5 TAAR5 vs. 2,284 ± 105 T5tg; n = 3 mice of each genotype). The number of glomeruli per bulb was similarly increased from ∼2 in *T5-RFP* mice^32^ to 36.2 ± 1.3 (mean ± SEM; n = 6 bulbs from 3 mice) in *T5tg* mice. The sensitivity of T5tg-expressing OSNs is indistinguishable from those expressing the native TAAR5 gene (Fig S6). Because TAAR5 is the most sensitive receptor for TMA^37^, the transgene dramatically increased the number of glomeruli responding to near-threshold concentrations of TMA (Fig S6). Thus, the transgene rescued aggressive behaviors despite a dramatic functional remapping of TAAR5 inputs.

## Discussion

Here, we report that sensory input via the main olfactory receptor TAAR5, responding to the male urinary odor TMA, mediates valence responses that are dependent on social status, and drives inter-male aggression in mice. Our experiments demonstrate that TMA is an aggression-promoting, male social cue, and identify a specific main olfactory input that exerts a strong influence on aggression in mice. These findings reveal a striking influence of a single main olfactory receptor on social behavior in mammals.

Urinary scent marks advertise the identity and social status of the marking male and prime the receiver for an appropriate behavioral response^48-50^. Male mouse urine has been reported to contain an “aversive pheromone” that is detected by the main olfactory system^51^. While chemicals have been isolated that may influence responses to urine^52,53^, the repertoire of urinary chemicals that act as an aversive/attractive cues is not completely characterized. Our data show that male behavioral responses to TMA parallel previously characterized responses to whole urinary scent marks^54-56^. Dominant males exhibit prolonged investigation of unfamiliar dominant male urine^48^, likely assessing potential competitors or intruders. In contrast, subordinate males avoid unfamiliar dominant male urine^54^, presumably in an attempt to reduce the likelihood of aggression. Thus, TAAR5 may play a crucial role in how male mice interpret scent marks.

Previous studies have reported that TMA is attractive to mice, similar to what we see for dominant males^39-41^. In the study of Li *et al*., mice were individually housed, and socially isolated males typically exhibit dominant behaviors^45^. The degree of attraction observed in other studies is likely influenced by methodological differences in addition to social status. In any event, our studies reveal, for the first time, a reliable, state-dependent shift in behavioral responses to a male-derived odor, suggesting that TAAR5 may influence social behavior more broadly.

In support of this, we show that TMA, acting through TAAR5, directly enhances aggression during male-male social interactions. Given that it is highly enriched in adult male mouse urine^38,39^, our data suggest that TMA is a key social odor that acts as an indicator of male identity and that is required to elicit full aggression in paired males.

Despite reduced aggression, TAAR5 deletion males still establish a social hierarchy. It is also important to note that while aggression is reduced in the TAAR5 deletion pairs, it is not absent. This residual aggression is likely dependent on other sensory cues^57,58^. Thus, while we have identified a novel main olfactory input that contributes to aggression, our data highlight that complex male social interactions involve multiple olfactory subsystems working in concert.

The TAARs are highly conserved across vertebrate evolution from fish to humans^59^. How the TAAR gene family contributes to olfactory function is still not completely understood. In mice, the TAARs are the most sensitive receptors for amines^37^—a function that is likely advantageous across phyla. Moreover, TAAR4 plays a role in interspecific communication by mediating behavioral avoidance of the predator odor, phenylethylamine^42^. We now show that TAAR5 influences inter-male aggression, identifying a previously unknown role for TAARs in a defined social behavior.

While there is great interest in olfactory receptors that mediate specific social behaviors^16-19^ few have been identified in the mammalian main olfactory system. Sensory neurons that express the receptor GC-D (a non-GPCR) and that mediate a defined behavior send axons to a subset of atypical (necklace) glomeruli in a distinct compartment of the olfactory bulb^28^. The large array of typical glomeruli in the main olfactory bulb is divided into receptor-specific glomerular domains ^32,34^ which have been proposed to relate to innate vs learned odor responses^34-36^. TAAR5-mediated aggression provides a unique opportunity to test the idea that input to specific glomeruli is critical for appropriate behavioral responses. In fact, we observed that the aggression-promoting function of TAAR5-persists despite gross remapping of TAAR5 glomeruli in the olfactory bulb. This could indicate that the behavioral responses to TMA is wholly learned, or that dedicated circuits downstream of the glomerular inputs rewire to compensate for the different receptor input pattern^41^. In any case, our data demonstrate a high degree of flexibility in the peripheral representations of a male-specific odor that influences an important social behavior.

## Acknowledgements

This work was supported by grants from NIH/NIDCD to TB (R01DC013576), AC (R21DC018905), and AD (F32DC012004 and R03DC014565). We thank David Swygart for collecting preliminary data, and Amanda Menzie for technical support. We acknowledge Rada Norinsky and the Transgenic Services Laboratory at The Rockefeller University for transgenic mouse production, and Gail Guth for artistic rendering of mouse ethograms.

## Author Contributions

A.D., A.C. and T.B conceived the project and designed the experiments. A.D., A.C., and K.B performed behavioral experiments. J.Z. performed electrophysiological recordings. A.C performed in vivo imaging experiments. S.K. and T.T. performed gene expression analysis. P.F. conceived the transgenic strategy for receptor overexpression and facilitated transgenic mouse production. A.C., A.D. and T.B. wrote the manuscript with input from all authors.

## Methods

All procedures were approved by the Northwestern University Institutional Animal Care and Use Committee.

### Mouse Strains

The *ΔTAAR5-LacZ (ΔT5*) mouse strain (*Taar5*^*tm1(KOMP)Vlcg*^) was obtained from the Knockout Mouse Project repository (www.komp.org). The *neo* selection cassette was removed from the targeted allele by crossing to *HPRT-Cre* (*Hprt1*^*tm1(cre)Mnn*^). All analyses were done using the Cre deleted allele in mice lacking *HPRT-Cre*. The TAAR1 knockout strain^60^ was graciously provided by Dr. Marius Hoener. The *ΔTAAR4-YFP* allele was previously described^32^.

*<u>TAAR5-RFPtg</u>*: Transgenic expression of TAAR5 was achieved using a transgene consisting of a 6.8kb fragment of the *Olfr151* (M71) locus that was modified by 1) insertion of a 5x homeodomain sequence^61^ placed 490 bp upstream of the transcription start site, and 2) by replacing the Olfr151 coding sequence with the coding sequence for mouse TAAR5 flanked by Asc I sites and preceded by a Kozak consensus sequence (*GGCGCGCCACCATG*). The coding sequence was followed by an internal ribosome entry site and TauCherry (a fusion of bovine tau and monomeric cherry fluorescent protein. The transgene was column purified and injected into C57BL/6J zygotes to obtain transgenic founders.

To control for background effects, all experiments were performed using littermates from heterozygous crosses.

### Gene Expression Analysis

RNA was extracted from tissue samples using RNeasy RNA isolation kit (Qiagen). Samples were treated with DNAse to eliminate genomic DNA contamination. RNA quality was determined using the 2100 Bioanalyzer (Agilent Technologies). An RNA integrity number (RIN) was calculated and only samples with an RIN of greater than 8 were selected for further analysis. Reverse transcription was performed using Superscript III Reverse Transcriptase (Invitrogen) with 2 μg of total RNA.

Gene expression in sample cDNA was measured by quantitative PCR according to MiQE guidelines^62^. Samples were amplified using a Bio-Rad CFX384 Touch™ Real-Time PCR Detection System using iQ™ SYBR® Green Supermix (Bio-rad). Cycling conditions were: 95°C for 3 minutes, followed by 40 cycles of 95°C for 15s and 59°C for 45s. Reactions were 10 μL, containing 5 μL of SYBR Green Supermix, 300 nM of each primer, and cDNA equivalent to 30 ng of total RNA. All reactions were run in quadruplicate. Melt curve analysis was performed at the end of each qPCR run to verify amplicon specificity. No-Template Controls (NTCs) were run for every primer set on every plate. Genomic DNA was not detected in no-reverse-transcription controls (NRTs). Expression of *Taar5* was measured against *Gusb* as a reference gene. Primer efficiency was measured by constructing standard curves across template concentration. All primers had measured efficiencies of 98-102%. Primer sequences were: Taar5 fwd, 5’ ACTGTTGACACACTAGTGGAC 3’; Taar5 rev, 5’ GATGGGATTACAGGCTGAATTG 3’ (105 bp); Gusb fwd, 5’ GCTGATCACCCACACCAAAG 3’; Gusb rev, 5’ CACAGATAACATCCACGTACG 3’ (107 bp).

### Behavior

Mice were housed in same-sex groups of 2-5 individuals in a reverse 12/12 hr light - dark cycle and provided with food and water *ad libitum*. For mice used in place preference assays, cages were changed daily to prevent adaptation to amines that are present in mouse urine.

#### Male Social Interactions and Dominance Assay

Wild-type and homozygous ΔT5 littermates (or homozygous ΔT5 littermates with or without the TAAR5 transgene) between 12-16 weeks of age were individually housed with an adult female for one week. To establish dominance, non-sibling males of the same genotype but different coat colors were placed in a new cage, and the first 30 minutes of their interactions were recorded with a digital video camera under low-intensity red light during the animals’ dark phase. Following this recording period, the pair was housed together for one week, after which the social status of each male was determined by differential scent marking^43^. Briefly, males were placed in an acrylic chamber (12×18×9 in) on either side of a wire mesh divider. The wire mesh allowed chemosensory, visual, and auditory communication but prevented direct contact. The bottom of the chamber was lined with Whatman filter paper. Males were allowed to scent mark on their half of the chamber for one hour. Mice were then returned to their home cage and the scent marks were stained with ninhydrin (Tritech Forensics) and photographed. Dominance was assessed based on the number and size of urine marks^43^. Male pairs with clear dominant/subordinate relationships were tested in the valence assay (below) and their initial behavioural interactions were analysed.

Male behavioural interactions were scored blind to genotype and dominance status. The frequency of four instantaneous behaviours (pinning, biting, shovelling, and tail rattling) were recorded, and the frequency and duration of eight other behaviours (grooming, anogenital investigation, facial investigation, mounting, freezing, chasing, fighting, and defensive upright posture) were recorded using Solomon Coder software (solomoncoder.com). Behaviours were classified as described in the Stanford mouse ethogram (mousebehavior.org).

For painting experiments, adult wild-type males were housed with an adult female for a week to establish a dominant phenotype. To record behavioural interactions, males were placed in a new cage together with a P18 wild-type male which was painted with either 100 ul water or TMA (5 mM, diluted in water) in the urogenital region. The first 30 min of interactions were recorded with a digital video camera under low-intensity red light during the animals’ dark phase and analysed as described above.

#### Odor Valence Assay

Odor valence was determined using a place preference assay^42^ consisting of three phases: handling (2 days), pre-trial (2 days), odor trials (2 days). Handling habituated the mice to the experimenter and consisted of individual mice resting in the experimenter’s open gloved hand for 2 minutes. The pre-trials and odor localization tasks were performed in clean, autoclaved 30 x18 x12 cm plastic cage bottoms. The pre-trials habituated the mice to the experimental apparatus/procedure and were identical to the experimental trials except no odorant was used. A disposable curtain divided the cage into an odorized and non-odorized compartments with minimal air transfer between compartments. Each mouse was introduced to the larger compartment and allowed to habituate for 3 minutes, after which a 3.5 cm petri dish containing 20 μl of water on Whatman paper was introduced into the odorized compartment. The top of the petri dish was perforated to allow odorants to escape but prevented direct contact with the stimulus. Mice were allowed to interact with the petri dish for 3 minutes. The petri dish was then removed and another identical petri dish containing 20 μl of an odorant on Whatman paper was added for 3 minutes. Mice were naïve to each stimulus and were tested only once for a given odor. All trials were video recorded and the mouse’s location was tracked using Limelight 3.0 software (Actimetrics). The aversion index was calculated as the difference between the time spent in the odorized chamber when an odor (or water) was present and the time spent in the odorized chamber when water was present. Data from mice that showed a very strong preference for either chamber (<10 s or >170 s of the trial in the odorized chamber) during the initial water trial were discarded. All experiments and analysis were performed blind to genotype and social status.

#### Odor stimuli

Monomolecular stimuli consisted of 2-phenylethylamine (CAS# 64-04-0; Sigma 128945), and trimethylamine (CAS# 75-50-3; Sigma 92262). All dilutions were made in distilled water and are expressed as percent by volume.

#### Awake in vivo imaging

Imaging was performed in head-fixed, wheel-running mice ^37,63^. Briefly, 2-5 month old mice that were either positive or negative for the T5-RFPtg and heterozygous for the OMP-GCaMP3 allele were implanted with an optical window and headbar over the olfactory bulbs. After recovery, fluorescence signals were recorded using a digital camera (NeuroCCD 256, Redshirt Imaging) on a wide-field fluorescence microscope. Response maps were obtained by subtracting a temporal average preceding the stimulus from a temporal average at the maximum response.

#### Electrophysiological recordings

The olfactory epithelium from P10-P14 mice was removed and kept in oxygenated bath solution (95% O2-5%CO2), containing (in mM) NaCl 124, KCl 3, MgSO4 1.3, CaCl2 2, NaHCO3 26, NaHPO4 1.25 and glucose 15 (pH 7.4). The knobs of OSNs were visualized and targeted for recording using an upright DIC microscope (Zeiss Axioskop 2 Plus) equipped with a CCD camera (SensiCam QE; Cooke Corporation) and a 40x water immersion objective. Patch pipettes were pulled from borosilicate glass with a P-97 horizontal puller (Sutter Instrument, Novato, CA), and fire-polished using a microforge (MF 83; Narishige, Japan). Electrophysiological recordings were controlled by an EPC-10 amplifier combined with Pulse software (HEKA, Germany). Perforated patch clamp was performed by including 260 µM amphotericin B in the recording pipette, which was filled with the following solution (in mM): KCl 70, KOH 53, methanesulfonic acid 82.3, EGTA 5, HEPES 10, and sucrose 70 (pH 7.2). The electrodes had tip resistances ranging from 8-10 MΩ when filled with internal solution.

Odorants were applied via a multi-barrel pipette placed 20 µm downstream of the cell. Stimuli were delivered using a pressure ejection system (PDES-02D, NPI Electronics, Germany). Odorants were dissolved in dimethyl sulfoxide (DMSO) and stored at −20°C. Odorant concentrations were further diluted in bath solution as necessary. The final concentration of DMSO was less than 0.1%. Analysis and curve fitting were performed using Igor Pro 5.01 (WaveMetrics, Inc., OR). Dose-response curves were fitted by the Hill equation, I =Imax/(1+(EC50/C)^n^), where I represents the peak of odorant-induced responses.

#### Statistical Analysis

Odor aversion data were analysed by univariate ANOVA, stratified by odor. Aversion index was skewed and contained both positive and negative values. We therefore shifted the outcome by a constant so that all values were greater than zero, then log transformed. We included interaction terms to determine whether state differences varied by genotype. A simplified model was considered if the 2-way interaction was not found significant at a 10% significance level. Pre-specified contrasts were performed to compare aversion index between states separately for each genotype using a Bonferroni procedure to correct for multiple comparisons.

Male-male interaction behaviors were analysed as follows. Descriptive statistics were compiled for duration and frequency of behaviors (shoveling, grooming, anogenital investigation, face-to-face investigation, mounting, freezing, chasing, fighting, biting, pinning, defensive upright, and rattling) by genotype and social status; mean ± standard deviation, median (Q1-Q3), minimum and maximum were calculated for all variables. Due to the strong correlation between frequency and duration, we chose only to model frequency of behaviour in order to minimize the chance of type 1 error. To determine the association between social status and frequency of behavior, we performed negative binomial regression with log-link to account for over dispersion. We hypothesized the association between social status and frequency of behavior would differ between genotypes, thus we included a genotype by social status interaction when statistically significant (relaxed p-value<0.10). We then estimated the association between social status and frequency of behavior for each genotype using least-squared means or marginal means. The original statistical analysis planned to adjust for mouse pairs, however the variance due to pair was negligible. Thus, the random intercept for pair was excluded. Analysis of TMA painting interactions were done in a similar manner looking for an association between painting and frequency of behavior.

Analyses were conducted in R version 4.0.3 (R-Core Team, Auckland, NZ) and assumed a two-sided, 5% level of significance unless otherwise specified.

**Supplementary Figure 1.**
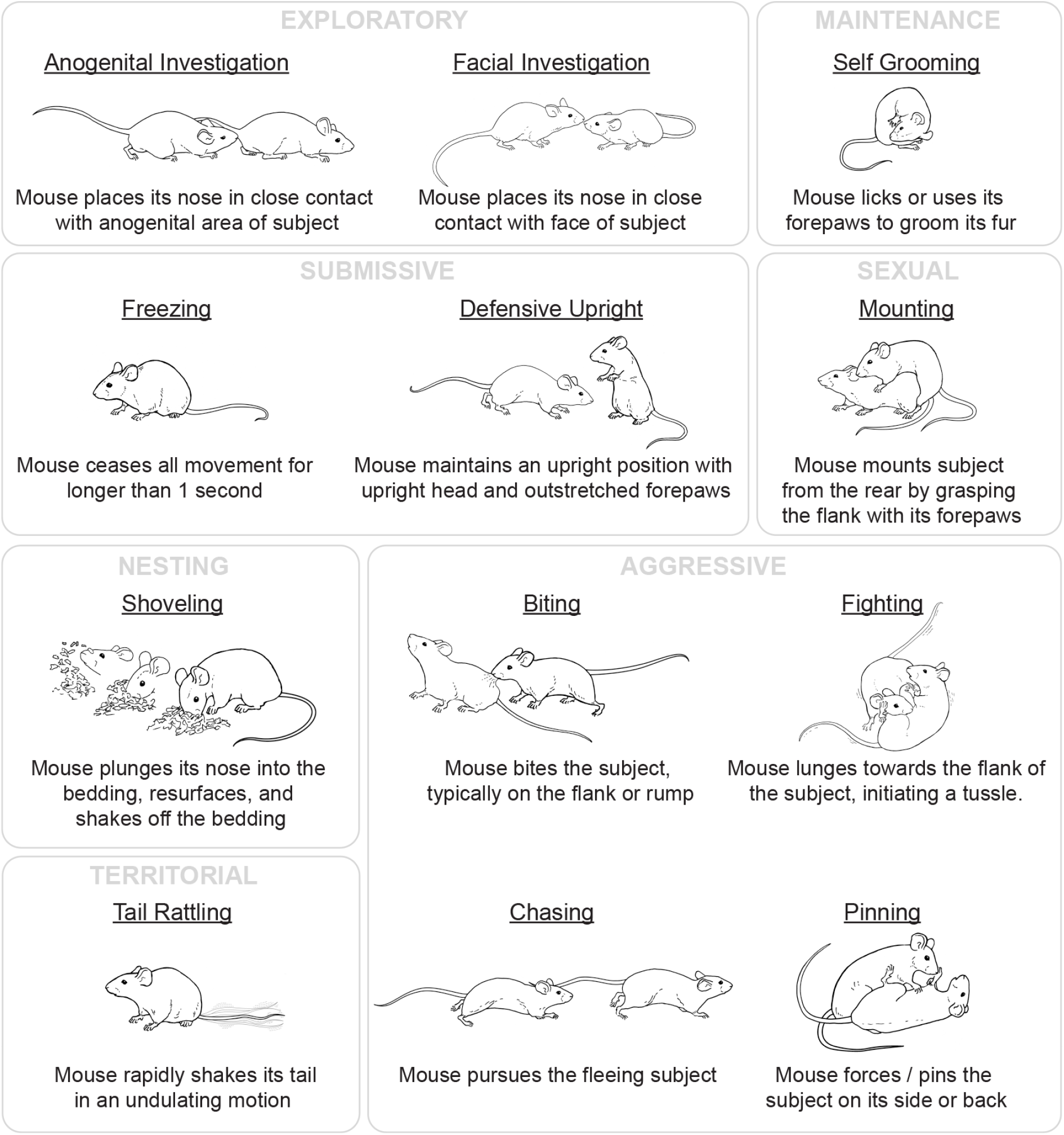
Ethogram of scored male-male behaviors. Behaviors are grouped by function, and the criteria for scoring each behavior are defined.

**Supplementary Figure 2.**
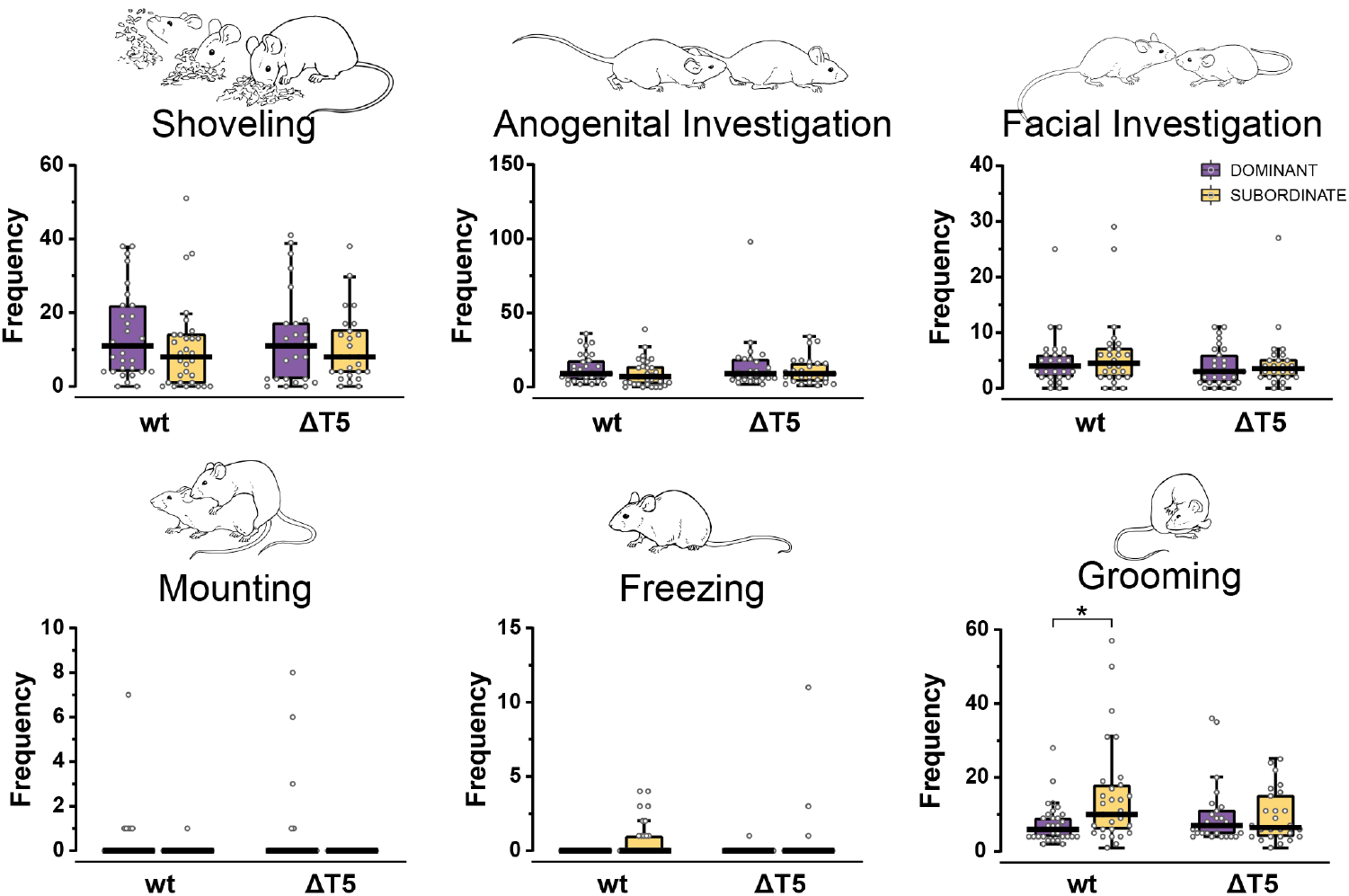
Quantification of non-aggressive behaviors in paired TAAR5 deletion and wild-type mice during 30 minute initial interaction. (Continuation of data shown in Fig 2). Behavior frequencies are plotted by genotype (wild-type vs. ΔTAAR5) and social status (dominant, purple vs. subordinate, yellow) based on which male initiated the behavior. (Thick lines, medians; boxes, 25th–75th percentiles; whiskers, most extreme data points not considered outliers, method of Tukey). n = 26-31 mice per experimental group. \**p* = 0.0001 (negative binomial regression).

**Supplementary Figure 3.**
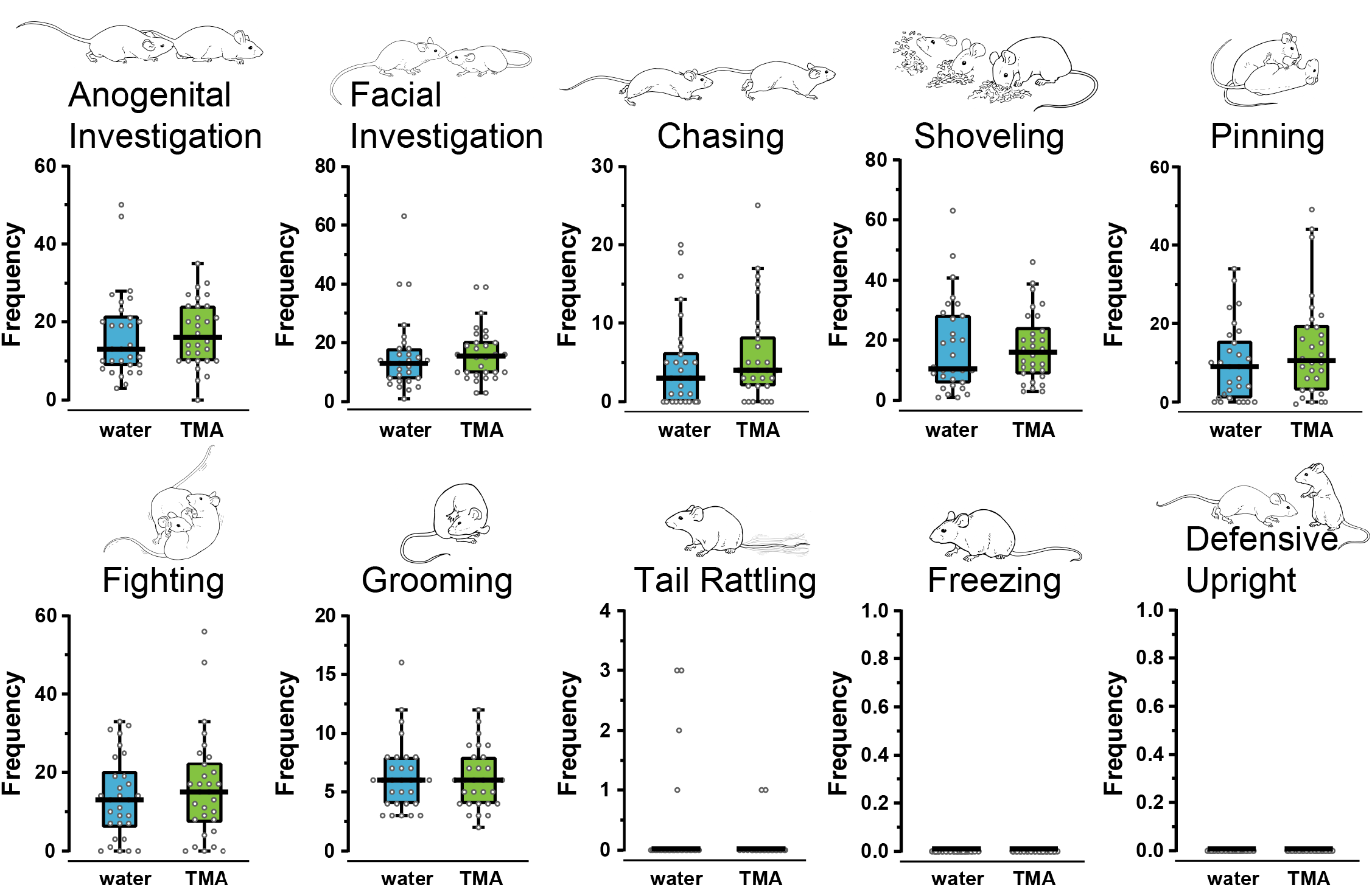
Quantification of behaviors in adult wild-type males paired with painted juvenile males (for behaviors not shown in Fig 3). Plots show frequencies for behaviors exhibited by the adult males during the first minutes, sorted by odor painted on the juvenile (water, blue or trimethylamine (TMA), green). Thick lines, medians; boxes, 25th–75th percentiles; whiskers, most extreme data points not considered outliers (method of Tukey).

**Supplementary Figure 4.**
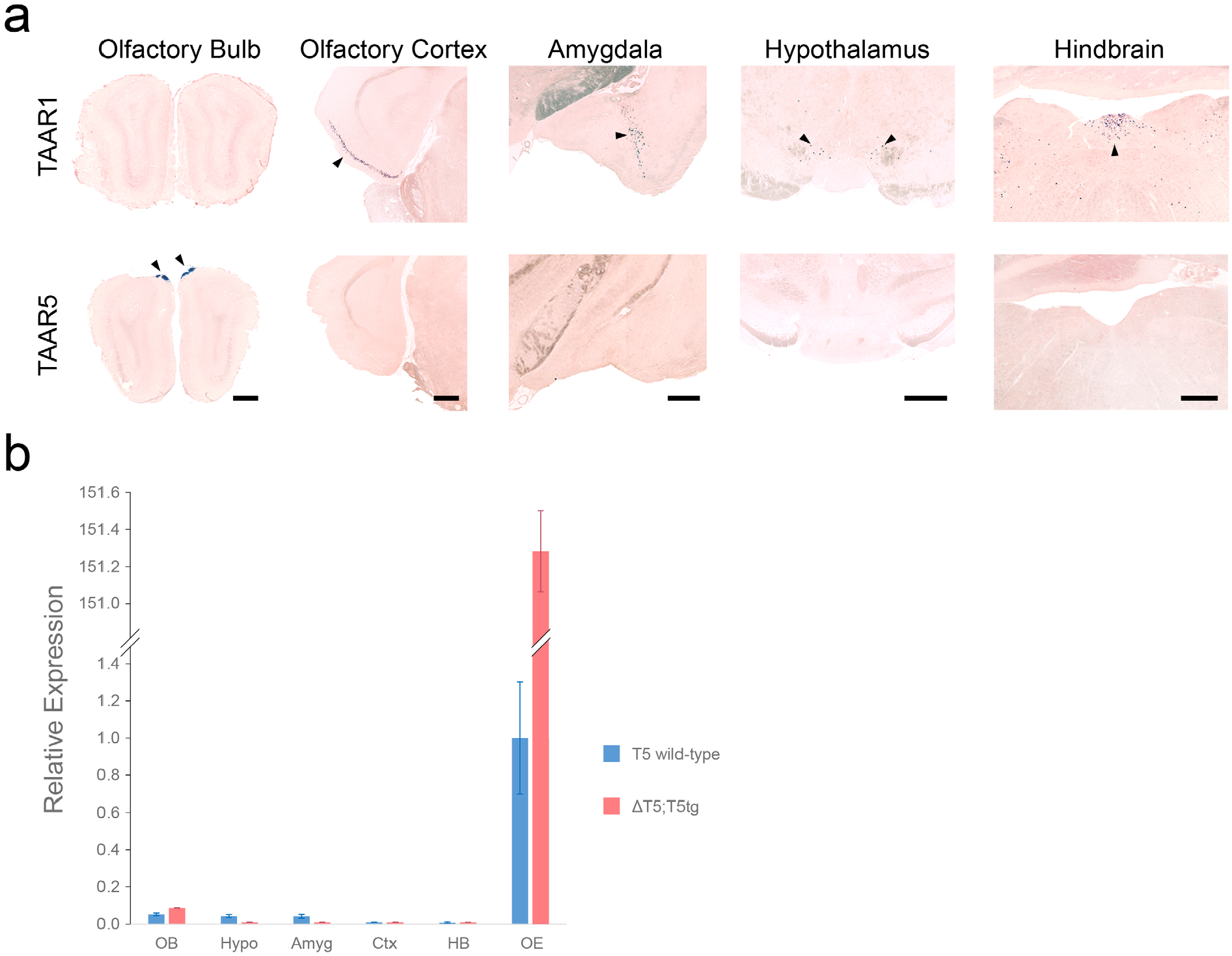
Selective expression of TAAR5 in the olfactory system. **a)** Histological sections of olfactory bulb, olfactory cortex, amygdala, hindbrain and hypothalamus, taken from TAAR5-LacZ and TAAR1-LacZ knock-out mice. *Taar1* is expressed at low levels in the brain^64,65^ and serves as a positive control. Sections were stained for β-galactosidase activity with X-gal (dark blue; arrowheads) for 24 hours and counterstained with neutral red. Expression of *Taar5* is observed only in the glomerular layer of the olfactory bulb. Expression of *Taar1* is observed in all other regions except the olfactory bulb. Data were obtained from 6 animals. Scale bars= 500 μm for each corresponding region. **b)** Expression of TAAR5 determined by qPCR from wild-type (n=7) and ΔTAAR5; T5tg (n=8) mice. Data are plotted relative to expression in wild-type olfactory epithelium. Break in the y-axis accommodates the high expression measured in the OE of transgenic mice. Inconsistent, low expression was observed in some hypothalamus and amygdala samples of wild-type mice. No expression was observed outside the olfactory system of ΔTAAR5; T5tg mice. Consistent expression in the OB of both genotypes is likely due to the small fraction of receptor mRNA in sensory axon terminals^66^.

**Supplementary Figure 5.**
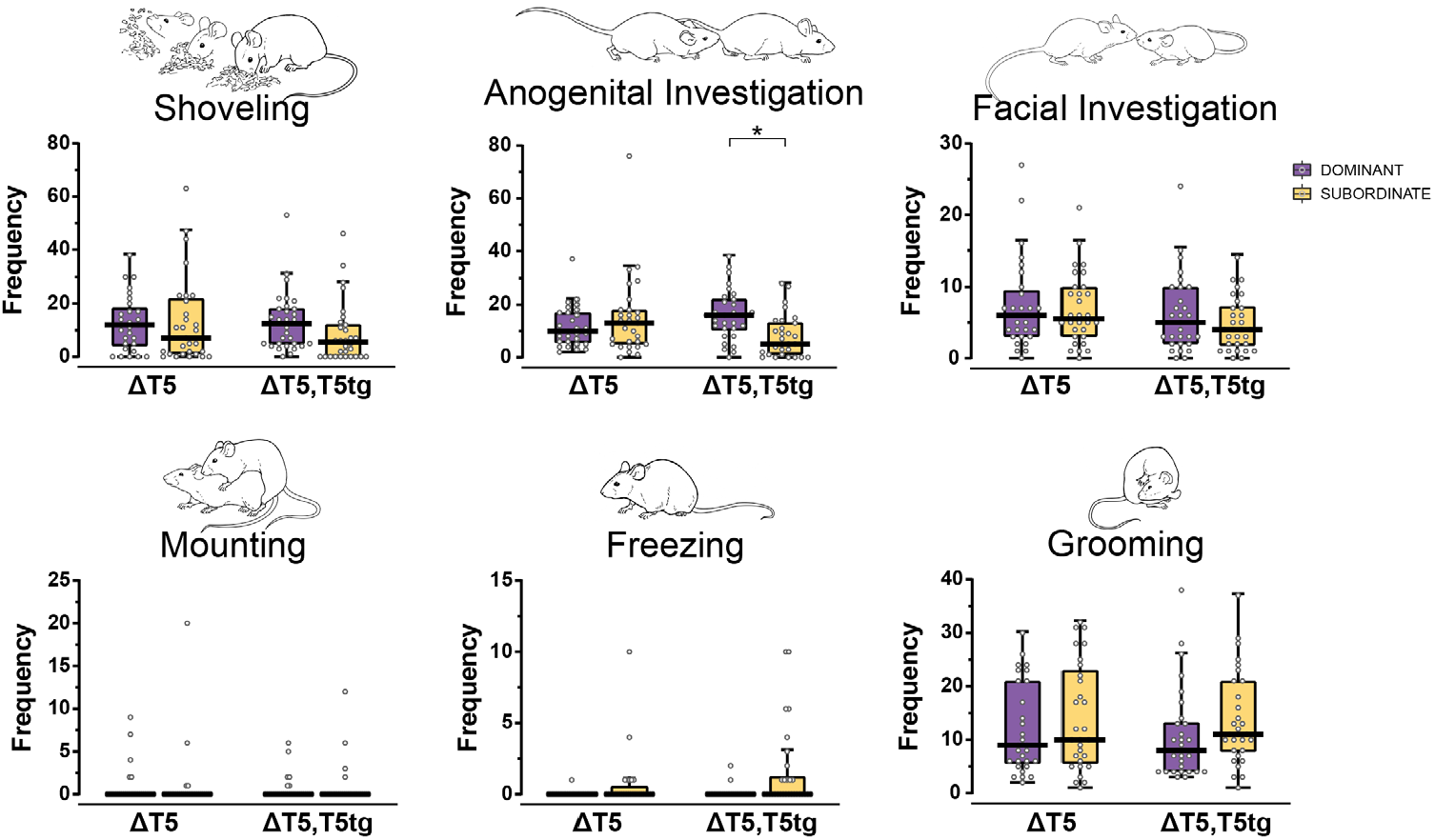
Quantification of assayed behaviors in paired TAAR5 deletion and TAAR5 transgenic rescue mice during 30 minutes of initial interaction. (Continuation of data shown in Fig 4). Data are sorted by genotype—ΔTAAR5 mice (ΔT5) and ΔTAAR5 mice that are hemizygous for the T5tg (ΔT5, T5tg)—as well as by social status (dominant, purple vs. subordinate, yellow). Thick lines denote medians; boxes, 25th–75th percentiles; whiskers, most extreme data points not considered outliers, method of Tukey). \**p* = 0.0011, anogenital investigation (negative binomial regression). n = 29-30 mice per experimental group.

**Supplementary Figure 6.**
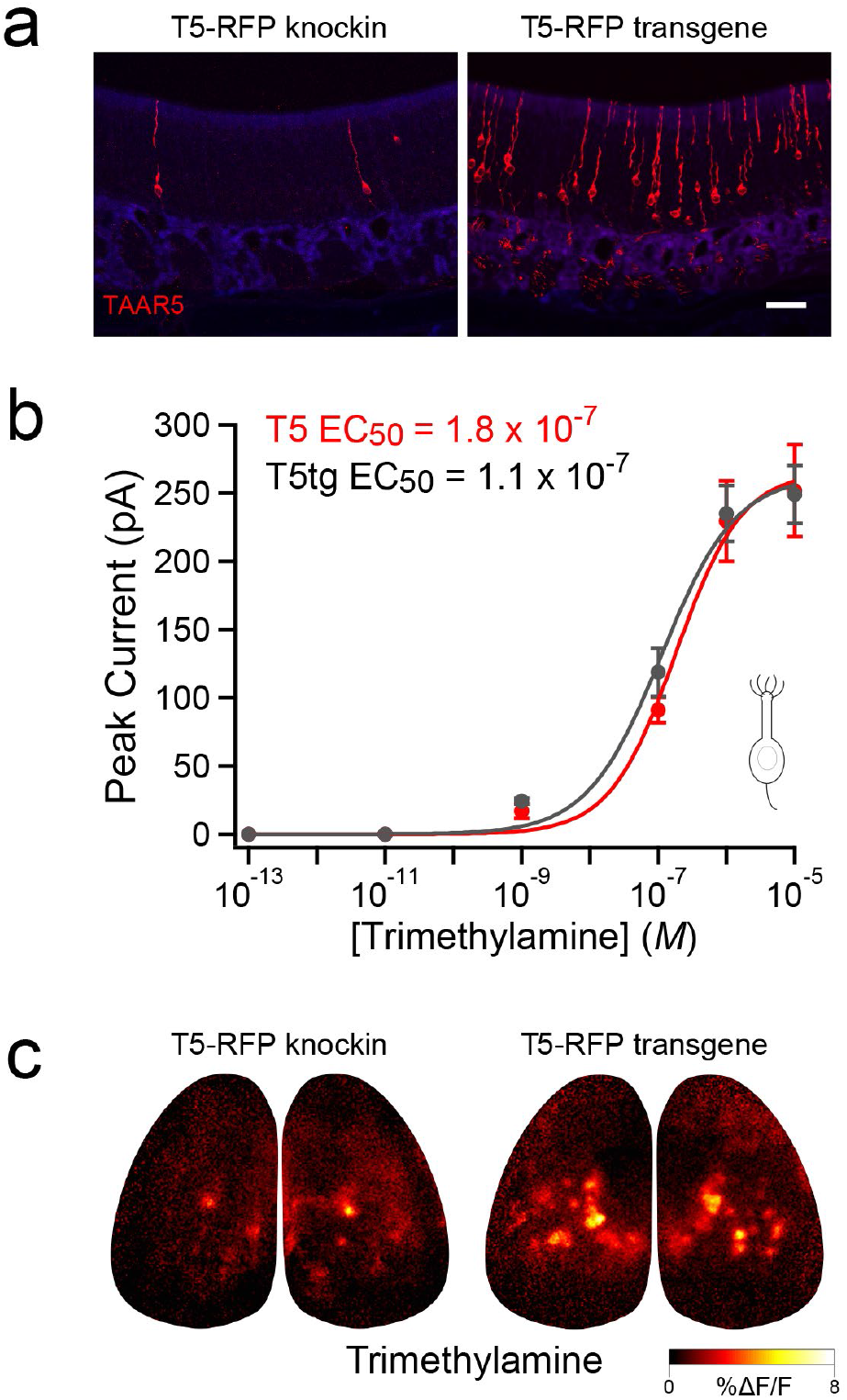
Transgenic expression of TAAR5. **a)** Confocal images showing expression of natively-tagged TAAR5 and (TAAR5-RFP knockin; red) and TAAR5-RFP transgene (T5-RFPtg, red) in OSNs of the olfactory epithelium. Scale bar = 40 μm. **b)** Dose-response data for trimethylamine measured via perforated patch clamp from OSNs expressing TAAR5-RFP (red; n=8 cells) and T5-RFPtg (black; n=7 cells). Peak response currents (mean ± SEM) are plotted. **c)** *In vivo* calcium imaging from OSN terminals in the olfactory bulb showing a greatly expanded spatial response to a near-threshold concentration of trimethylamine across glomeruli.

